# Diet and feeding strategies of two sympatric mouse lemurs (*Microcebus*) in the xeric forests of Andohahela, southeastern Madagascar

**DOI:** 10.64898/2026.08.22.746410

**Authors:** Sam Hyde Roberts, J. Carolina Segami, Vatosoa J. N. Harinala, Anne D. Yoder

## Abstract

Understanding how closely related species coexist in highly seasonal and unpredictable environments is central to studies of ecological differentiation and niche partitioning. We investigated the feeding ecology of sympatric populations of *Microcebus murinus* and *M. griseorufus* within a contact zone in Andohahela National Park, southeastern Madagascar, across dry and wet seasons. Using a combination of direct behavioral observations (1,611 feeding records), fecal sample analyses (n = 56), and vegetation phenology surveys, we quantified dietary composition, seasonal shifts in resource use, and habitat-related variation. Seasonal changes in diet were pronounced, with dry-season feeding dominated by exudates and wet-season diets incorporating greater proportions of fruit and flowers, closely tracking phenological patterns at both sites. Diets of both species were dominated by plant resources, but consistent interspecific differences in dietary strategy were evident. Although both species consumed comparable proportions of insect prey, *M. murinus* showed pronounced wet-season increases in the use of high-sugar, carbohydrate-rich floral resources (15.9%) and hemipteran-associated honeydew (30.6%). In contrast, *M. griseorufus* relied more consistently on predictable exudates throughout the year. Fecal analyses supported observational data but revealed differences in the detectability of dietary components, with increased representation of invertebrates in the wet season and seeds in the dry season. These results indicate substantial dietary overlap but consistent differences in resource use, suggesting that coexistence is facilitated by fine-scale trophic differentiation within a broadly shared omnivorous niche. Such subtle but persistent differences in feeding strategy likely reduce competitive overlap and enable continued sympatry in a climatically variable and resource-limited system.

**Graphical Abstract:** 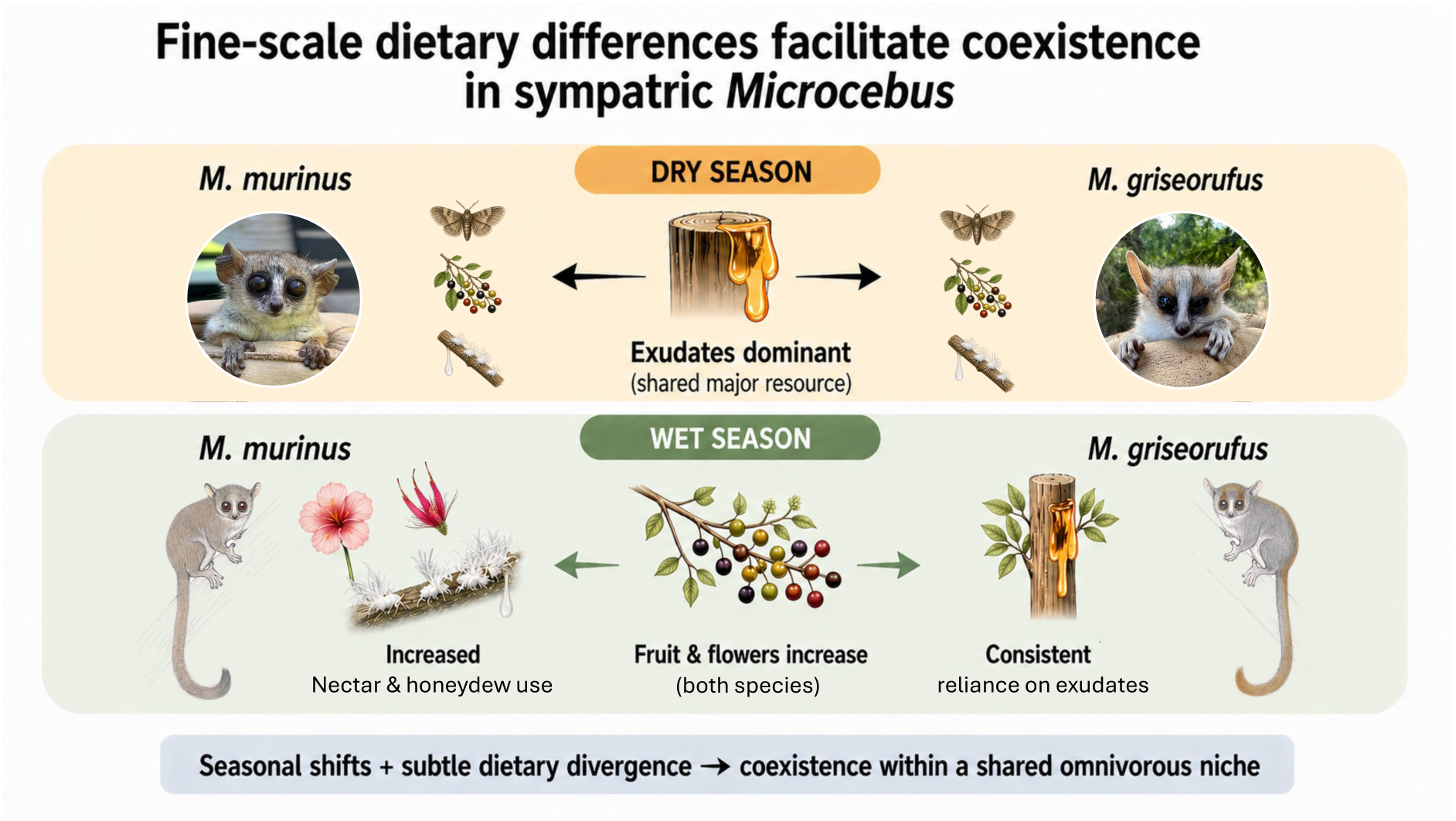

## Introduction

Understanding how closely related species coexist in shared environments remains a central question in ecology, particularly in systems where resources are limited, temporally variable, or spatially heterogeneous. Classical niche theory predicts that stable coexistence requires differentiation in ecological niches, thereby reducing interspecific competition for limiting resources (Gause, 1934; Hardin, 1960; Chase & Leibold, 2003; Levine & HilleRisLambers, 2009). In practice, such differentiation may arise across multiple, non-mutually exclusive axes, including spatial segregation, dietary partitioning, temporal differences in resource use, or variation in resource exploitation. Across a wide range of animal taxa, feeding niche differentiation is frequently identified as a primary axis structuring coexistence (Schoener, 1974; Ross, 1986; Van Holstein et al., 2024), particularly among ecologically similar, co-occurring species. Understanding these processes is therefore critical for predicting species persistence amid rapid environmental change and forecasted biodiversity losses (Barnosky et al., 2011; Estrada et al., 2017).

In primates, niche differentiation is a key mechanism facilitating coexistence among species and structuring community composition (MacKinnon & MacKinnon, 1980; Ganzhorn, 1989; Van Holstein et al., 2024). It operates across multiple scales, including functional traits such as body size and broad dietary strategies (Fleagle, 2013), as well as across ecological axes such as spatial and temporal patterns of resource use (Schreier et al., 2009). Together, these dimensions support the coexistence of complex, species-rich assemblages across many tropical systems (Estrada et al., 2017), with broader patterns of primate diversity closely linked to environmental productivity and resource availability (Kay et al., 1997). However, among closely related or ecologically similar species—particularly those sharing similar body size and dietary guild—dietary differentiation often represents the primary axis of niche separation (Houle, 1997; Schreier et al., 2009). The extent and form of dietary separation, however, vary markedly among systems, indicating that the role of diet in facilitating coexistence is strongly context-dependent. Niche differentiation is therefore not fixed, but varies with environmental conditions.

Mouse lemurs (*Microcebus* spp.) represent a diverse radiation of small-bodied nocturnal primates endemic to Madagascar and provide an excellent system for examining these dynamics. Early studies suggested that small primates may rely heavily on energetically dense animal prey to meet high metabolic demands (Hladik, 1979; Kay, 1984), yet mouse lemurs are now recognized as flexible omnivores with highly variable diets (Atsalis, 1999; Lahann, 2007; Crowley et al., 2014). These diets comprise a mixture of plant-derived foods—including fruit, nectar, flowers, and exudates—supplemented by invertebrates, insect secretions, and, infrequently, small vertebrates (Radespiel, 2006; Dammhahn & Kappeler, 2008a). This dietary flexibility enables them to exploit a wide range of habitats, from humid rainforests to xeric spiny thicket systems, and underpins their broad distribution across Madagascar. However, this pattern is not uniform across the genus, and some *Microcebus* species exhibit highly restricted distributions and show more limited ecological breadth (e.g., Knoop et al., 2018; Giertz et al., 2026).

Within the genus, multiple species occur in sympatry (e.g., Schmid & Kappeler, 1994; Zimmermann et al., 1998; Rasoloarison et al., 2000; Gligor et al., 2009; Poelstra et al., 2021), and an increasing number of contact zones between species are recognized. Notably, *Microcebus murinus* has one of the broadest geographic distributions of any mouse lemur species and occurs in sympatry with congeners across western and southern Madagascar, providing a valuable comparative framework for examining how ecological conditions influence patterns of niche differentiation. Mouse lemur species are largely cryptic in terms of gross morphology and broad ecological traits (Yoder et al., 2016), suggesting a high potential for interspecific competition; however, stable coexistence appears relatively common. Despite their crypsis, fine-scale ecological differentiation is evident on close inspection, including variation in social organization, sleeping site ecology, and dietary strategies (Radespiel et al., 2003; Dammhahn & Kappeler, 2008a,b; Thorén et al., 2011; Hyde Roberts et al., 2026a,b). Across Madagascar, mouse lemurs also commonly occur in sympatry with other cheirogaleids (e.g., *Cheirogaleus* spp.), where dietary differences again appear minimal, with coexistence instead associated with temporal, spatial, or microhabitat segregation (Lahann, 2007). The widespread occurrence of sympatric lemur species across a diverse array of environmental contexts thus provides a powerful framework for testing how ecological conditions shape coexistence. Despite this, the processes facilitating coexistence among sympatric mouse lemur species remain incompletely understood.

Despite growing interest in mouse lemur dietary ecology, relatively few species have been studied in detail, reflecting both logistical challenges and the rapid increase in recognized diversity. As a result, our understanding of dietary variation across the genus remains limited (Hladik et al., 1980; Atsalis, 1999; Génin, 2001; Dammhahn & Kappeler, 2008a). In the few systems examined to date, subtle differences in the relative importance of key dietary resources, particularly in relation to seasonal phenological dynamics, appear to play an important role (Dammhahn & Kappeler, 2008a; Thorén et al., 2011). Thus, dietary dynamics appear to play a central role in niche differentiation, in concert with other ecological axes (Hyde Roberts et al., 2026b), highlighting both the ecological flexibility of the genus and the importance of environmental context in shaping feeding strategies.

Habitat and climate play important roles in structuring dietary strategies in mouse lemurs along environmental gradients. In humid rainforest systems, where fruit resources are relatively abundant and predictable, diets are dominated by fruit and other ephemeral plant resources (Atsalis, 1999). In contrast, species inhabiting drier western and southern forests rely more heavily on exudates and other stable food sources, reflecting increased seasonality and reduced predictability of fruit availability (Génin, 2001, 2008). In these environments, insect-derived secretions, such as honeydew produced by sap-feeding hemipterans, also represent important supplementary carbohydrate resources (Corbin & Schmid, 1995; Dammhahn & Kappeler, 2008a; Thorén et al., 2011). Dietary strategies therefore align along a gradient from temporally variable, high-reward resources to more predictable, low-variance resources. However, these patterns are further shaped by interactions among sympatric species, such that observed dietary strategies reflect both environmental constraints and the outcomes of niche differentiation within communities. Together, these patterns indicate that the mechanisms underlying coexistence are shaped by the availability, distribution, and predictability of key resources.

In Madagascar, environmental seasonality is often coupled with high climatic unpredictability, particularly in southern regions (Dewar & Richard, 2007; Génin, 2008). Under these conditions, resource availability varies both seasonally and between years, limiting the ability of individuals to anticipate periods of scarcity. Such environments are expected to favor either highly flexible foraging strategies or increased reliance on stable, predictable resources that buffer against environmental variability. Here, we investigate the feeding ecology of sympatric populations of *Microcebus murinus* and *M. griseorufus* within a contact zone in Andohahela National Park, southeastern Madagascar (Gligor et al, 2009; Rakotondranary et al., 2011; Rakotondranary & Ganzhorn, 2012; Hyde Roberts et al., 2026a). This region is characterized by pronounced seasonality and strong habitat heterogeneity, including spiny thicket and transitional forest systems (Andriaharimalala et al., 2011; Goodman et al., 2018). This system provides a direct test of how environmental context— particularly resource predictability—influences niche differentiation among sympatric species. Elsewhere, *Microcebus griseorufus* inhabits some of the driest and least predictable habitats in Madagascar and relies heavily on exudates and other stable resources (Génin, 2008), suggesting a close link between its feeding ecology and environmental unpredictability. In contrast, *M. murinus* is widely regarded as a flexible generalist, capable of exploiting a broad range of food resources across diverse habitats. However, direct comparisons between sympatric populations of these species under highly seasonal and unpredictable conditions remain lacking, and it is therefore unclear whether coexistence is structured primarily by differences in resource use or broader strategies linked to environmental predictability.

We combine direct behavioral observations, fecal analyses, and vegetation phenology surveys to quantify dietary composition and assess seasonal and habitat-related variation in resource use. Specifically, we test whether sympatric *M. murinus* and *M. griseorufus* (1) differ in their use of major food categories and plant resources, (2) exhibit seasonal shifts in diet corresponding to variation in resource availability, and (3) show evidence of dietary differentiation that may facilitate coexistence. By integrating multiple lines of evidence, we aim to identify the ecological mechanisms underlying coexistence and evaluate the extent to which differences in resource-use strategies—particularly along gradients of resource predictability—contribute to niche differentiation in highly seasonal environments.

## Methods

### Ethics statement

All field procedures involving animals were conducted in accordance with national and international guidelines for the ethical treatment of wildlife. All methods adhered to the principles for the ethical treatment of non-human primates and the Code of Best Practices for Field Primatology established by the American Society of Primatologists and complied with the legal requirements of Madagascar. Field research was conducted under permits granted by the Malagasy Ministry of Environment and Sustainable Development through the Direction des Aires Protégées des Ressources Naturelles Renouvelables et des Ecosystèmes (096/24/MEDD/SG/DGGE/DAPARNE/SCBE.Re, issued 27 March 2024; and 341/24/MEDD/SG/DGGE/DAPARNE/SCBE.Re, issued 13 September 2024). All animal handling and sampling procedures were approved by the Duke University Institutional Animal Care and Use Committee (IACUC; protocol no. A163-22-09) and were designed to minimize stress and disturbance. The use of radio collars and tissue sampling was justified by the need to investigate ecological and behavioural processes, and all individuals were released at their site of capture following processing.

### Study area

Fieldwork was conducted in southeastern Madagascar, with a primary focus on a verified contact zone between *Microcebus murinus* and *M. griseorufus* in Andohahela National Park (Gligor et al., 2009; Hapke et al., 2011; Hyde Roberts et al., 2026a) (Figure 1). Andohahela National Park (76,269 ha) comprises three noncontiguous parcels spanning a pronounced ecological gradient from humid evergreen forest to dry spiny thicket. Our study focused on Parcel II, which forms a sharp ecotone between transitional forest and spiny thicket habitats. Parcel II is characterized by a sub-arid climate with highly seasonal rainfall (<1000 mm annually), a cool dry season from June to August, and peak temperatures between September and November (Goodman et al., 2018). Vegetation is sparse and xerophytic, with strong seasonal limitations on plant productivity. Prolonged droughts and high inter-annual climatic variability make resource availability unpredictable, providing a natural setting for examining feeding strategies of sympatric species in a climatically extreme environment. Two study sites within Parcel II, Mangatsiaka and Tsimelahy, located approximately 10 km apart, were used for intensive behavioural observations. Tsimelahy (24.945° S, 46.630° E; 168 m a.s.l.) represents transitional forest, whereas Mangatsiaka (24.964° S, 46.555° E; 95 m a.s.l.) is characterized by dry spiny thicket vegetation.

**Figure 1.**
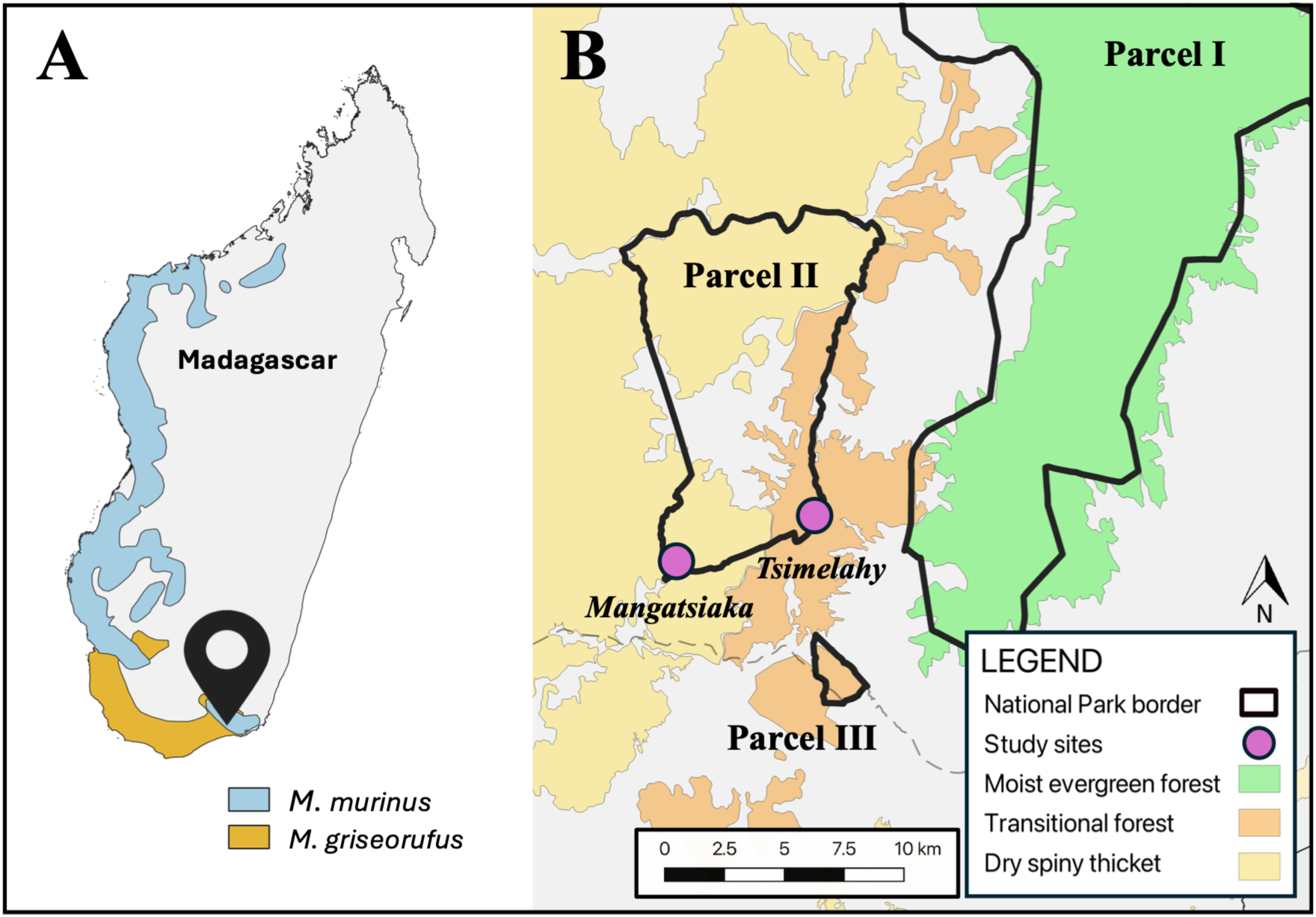
Geographic distribution of the study species and location of study sites in southeastern Madagascar. (A) Distribution ranges of *Microcebus murinus* and *M. griseorufus* across Madagascar. (B) Location of study sites, Mangatsiaka and Tsimelahy, within Andohahela National Park (Parcel II), in relation to major habitat types.

### Capture, handling, and species identification

Mouse lemurs were captured by hand during two field seasons between May–July 2024 and October–December 2024. A total of 104 individuals were captured at Mangatsiaka and Tsimelahy. Animals were sexed, weighed, assigned a unique identifier, and temporarily anesthetized with ketamine (10 mg/kg^-1^; 50 mg mL⁻¹ solution) for morphometric measurements and tissue sampling. A small ear-pinna biopsy was collected and preserved in ethanol for genetic analyses. Animals were monitored during recovery and released at their capture location. Species identity was validated using whole-genome sequencing and population genetic clustering analyses. Species assignment for each cluster was confirmed using mitochondrial cytochrome *b* sequences compared with reference databases. Characteristic phenotypes are presented in Figure 2 (panel D).

**Figure 2.**
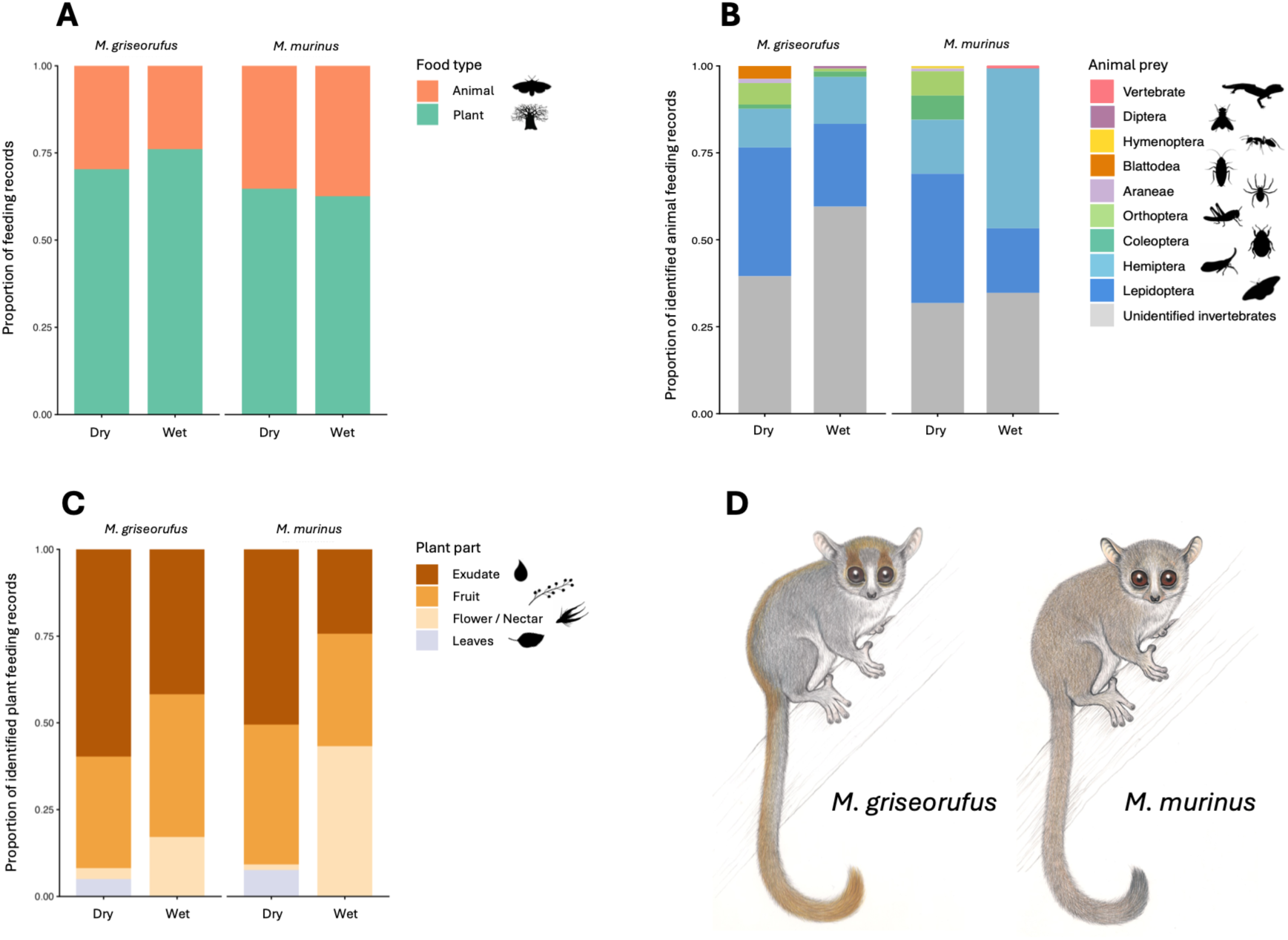
Dietary composition and feeding ecology of sympatric mouse lemurs at Andohahela National Park. (A) Relative contribution of plant versus animal feeding records across dry and wet seasons. (B) Composition of animal-derived feeding, showing major prey categories and hemipteran-associated honeydew feeding (Flatidae). Hemiptera records represent honeydew feeding, except for a single cicada prey record (*M. griseorufus*, Tsimelahy, wet season). (C) Use of plant parts (exudates, fruit, flowers/nectar, leaves) based on identified feeding records. *Leaves* refers to leaf-associated feeding events, likely including surface feeding or the use of plant- or insect-derived secretions. (D) Representative phenotypes of the study species, *Microcebus griseorufus* and *M. murinus*.

### Telemetry and behavioral observations

56 adult individuals were fitted with lightweight VHF radio collars (ATS M1420; ∼1.5 g). Collars were deployed across species, sexes, and sites during both dry and wet seasons. Animals were followed three to four times per week until battery life necessitated collar removal. Each focal animal was followed from dusk to dawn using standard radio-telemetry techniques. Behavioral data were collected using instantaneous scan sampling at 10-minute intervals, supplemented by continuous observations when visibility allowed (Altmann, 1974). Feeding events were recorded whenever observed, including plant or prey type and feeding behaviour where possible. A total of 224 full-night follows were conducted, representing 2,464 hours of observation.

### Feeding data standardization

Feeding records were standardized prior to analysis. Each observation was assigned a plant or insect taxon based on local vernacular names, field identification and available published resources (Schatz, 2001). When plant parts could not be confidently identified, items were classified as “unknown plant part” but retained for analyses of plant taxon use. Food items were grouped into functional categories: fruit, flower/nectar, exudate, leaf (plant) or vertebrate and invertebrate (animal) prey. Exudates included plant-derived gum and sap. Hemipteran-associated feeding (e.g., honeydew secretions) was treated separately from plant exudates and classified as an animal-derived resource. Invertebrate records were standardized across vernacular descriptors. Observations that could not be confidently classified beyond these levels were retained using conservative categories.

### Fecal sample collection and analysis

Fecal samples were collected opportunistically during the handling phase of captured individuals. A total of 56 samples were obtained, comprising 28 samples from the dry season and 28 from the wet season. Dietary remains were identified to the lowest possible taxonomic or functional level, including insect fragments, plant tissues, seeds, and other recognizable material. Items were classified into the same functional dietary categories used for behavioral observations to allow qualitative comparison between observational and fecal datasets. Fecal analyses were used to complement feeding observations and to verify the presence of dietary components that may be underrepresented in direct behavioral records.

### Habitat and vegetation surveys

Vegetation surveys were conducted at both study sites using 1-km transects (10 m width). All trees ≥15 cm in circumference were recorded for species identity, height, and diameter. Phenological status (leafing, flowering, fruiting) was assessed during both seasons to capture seasonal variation in resource availability. Tree community diversity and structure were quantified using standard diversity indices. Species accumulation curves and richness estimators were used to assess sampling completeness. Phenological status was standardized into four primary botanical states: flowering, fruiting, new leaves, and inactive. Leaf shedding, senescent foliage, and other non-productive leaf conditions were classified as inactive for feeding-ecology analyses, as these states do not represent resource-producing phases.

### Statistical analyses

Feeding records were classified according to food type (plant or animal), plant taxon, and, where possible, plant feeding category (fruit, flower, exudate, or leaves). Analyses of plant feeding categories were restricted to feeding observations for which the plant part consumed could be reliably identified. Feeding records were summarized as proportions of food categories, plant taxa, and plant feeding categories for each species, site, and season; percentages are presented as proportions of feeding records. Overall trophic composition (plant versus animal feeding) was quantified separately for each species, site, and season. Because hemipteran-associated feeding represents an animal-derived sugar resource that is functionally distinct from plant exudates, trophic composition was evaluated both including and excluding these records to distinguish between plant-derived resources and animal prey. Differences in trophic proportions between seasons within species, and between species within seasons, were evaluated using Pearson’s chi-squared tests. To test for overall differences in trophic composition, including potential interactions between species and season, binomial generalized linear models (GLMs) with a logit link function were fitted with feeding category (animal versus plant) as the response variable and species, season, and their interaction as predictors.

Variation in plant feeding categories was analyzed using multinomial logistic regression models with plant feeding category as the response variable and species, season, and site as predictor variables. Model significance was assessed using likelihood-ratio tests comparing nested models. This allowed evaluation of independent and interacting effects of species identity, seasonal variation, and habitat type on plant-part selection. Dietary composition based on plant taxa was summarized descriptively to quantify dietary richness, proportional use of plant taxa, and seasonal shifts in dominant food species. All statistical analyses were conducted in R (version 2026.01.0+392) using the packages dplyr, tidyr, ggplot2, vegan, and nnet, with significance assessed at α = 0.05.

## Results

### Dietary composition

A total of 1,611 feeding records were analyzed across both species, sites, and seasons (Table 1). Diets of both mouse lemur species were dominated by plant resources, although clear interspecific differences were evident in trophic composition (Figure 2). Across all observations, plant items accounted for 74.1% of feeding records in *Microcebus griseorufus* and 63.6% in *M. murinus*, with animal resources comprising 25.9% and 36.4% of records, respectively. Seasonal patterns were broadly similar between species, although *M. murinus* consistently consumed a greater proportion of animal-derived resources in both seasons than *M. griseorufus*. Across sites, the overall balance between plant resources and animal resources remained broadly similar, but the relative contribution of specific resources differed strongly, particularly with respect to hemipteran-associated feeding, floral resources, and plant exudates. Together, these results indicate broadly similar trophic structure between sites and species, but clear interspecific and habitat-related differences in the types of resources used within that shared framework.

**Table 1.** Seasonal and site-specific variation in dietary composition of sympatric *Microcebus griseorufus* and *M. murinus* at Andohahela National Park.

| Species | Site | Season | Total records (n) | Plant n (%) | Animal n (%) |
| --- | --- | --- | --- | --- | --- |
| *M. griseorufus* | Mangatsiaka | Dry | 143 | 106 (74.1%) | 37 (25.9%) |
|  |  | Wet | 212 | 140 (66.0%) | 72 (34.0%) |
|  | Tsimelahy | Dry | 130 | 86 (66.2%) | 44 (33.8%) |
|  |  | Wet | 314 | 260 (82.8%) | 54 (17.2%) |
| *M. murinus* | Mangatsiaka | Dry | 147 | 94 (63.9%) | 53 (36.1%) |
|  |  | Wet | 182 | 123 (67.6%) | 59 (32.4%) |
|  | Tsimelahy | Dry | 219 | 143 (65.3%) | 76 (34.7%) |
|  |  | Wet | 264 | 156 (59.1%) | 108 (40.9%) |

Plant resources include fruits, flowers, and exudates (tree gums and nectar), while animal resources comprise vertebrate and invertebrate prey, as well as insect (Hemiptera) secretions. Percentages are calculated relative to the total number of feeding records within each species x site x season combination.

Plant feeding involved a total of 81 identified plant taxa across both species, sites, and seasons. Dietary plant richness was broadly similar between species, with 62 taxa recorded for *M. griseorufus* and 54 taxa for *M. murinus*, indicating substantial interspecific overlap in plant resource use. Plant taxon richness was also comparable between sites (Mangatsiaka: 51 taxa; Tsimelahy: 49 taxa) and between seasons (dry: 53 taxa; wet: 49 taxa), suggesting that seasonal diets relied on broadly similar sets of plant taxa. Complete lists of plant food taxa recorded for each species are provided in Supplementary Tables S1A–S1B.

Specific plant feeding categories were identified for 600 of 1,108 plant feeding records (54.1%), and analyses of plant food types were therefore restricted to this subset. Exudates constituted the most frequently consumed plant resource for both species but were proportionally more important in the *M. griseorufus* diet (51.1%) than in the *M. murinus* (40.7%) diet. Fruit consumption was similar between species (36–37% of identified plant feeding records), whereas *M. murinus* consumed flowers and nectar more frequently (17.3%) than *M. griseorufus* (9.8%). Leaf feeding represented only a minor component of the diet in both species (≤5%) and appeared only during the dry season. Records reflect leaf-associated surface feeding, likely including plant- or insect-derived residues, rather than true folivory. Overall, plant-part use differed significantly between species (Pearson’s χ² = 11.62, df = 3, *p* = 0.009), indicating interspecific differences in the relative importance of exudates and floral resources despite broadly similar levels of fruit consumption.

Exploitation of specific plant parts was concentrated on a limited number of key plant taxa. Exudate feeding was strongly dominated by Taly (*Terminalia seyrigii*; Combretaceae), which accounted for 60.5% of all exudate feeding records, followed by Daro (*Commiphora aprevalii*; Burseraceae) (9.1%) and Jahiby (*Operculycaria decaryi*; Anacardiaceae) (8.7%), with all other taxa contributing comparatively minor proportions (<5% each). Floral feeding was similarly concentrated, with flowers of Varamomange (*Cloiselia carbonaria*; Fabaceae) comprising 54.3% of all flower-feeding observations, while Maramdoha (*Albizia* sp.; Fabaceae; 8.6%), Manongy (*Zanthoxylum* sp.; Rutaceae; 7.4%), and Hazonta (*Rhigozum madagascariense*; Bignoniaceae; 6.2%) represented secondary floral resources. These dominant plant-part resources were used by both species, although exudate feeding contributed proportionally more to the diet of *M. griseorufus*, whereas floral feeding was more prominent in *M. murinus*. Together, these results indicate that although a wide diversity of plant taxa was consumed overall, exploitation of specific plant parts—particularly gums and flowers—was strongly structured around a small number of key resource species.

Among animal-derived feeding records (n=503), unidentified invertebrates comprised the majority of observations (40.7%). Identifiable prey records (n=176) were dominated by moths (79.0%) while all other prey categories—including beetles (6.8%), crickets (5.1%), grasshoppers (3.4%), cockroaches (1.7%), and spiders (1.1%)—contributed only minor proportions. Rare additional observations included a large dipteran, a single hymenopteran (ant), and ingestion of caterpillar frass. Moth prey consisted predominantly of small, pale individuals (likely Microlepidoptera), with occasional larger specimens identified as hawk moths (Sphingidae) and silk moths (Saturniidae). Hemipteran-associated feeding, primarily involving honeydew from flatid planthoppers (*Flatidia* sp., Flatidae), comprised a substantial component of total animal-derived feeding (24.3% of records). These observations largely reflect sugar feeding rather than active prey capture, as individuals were frequently observed licking plant surfaces or planthopper aggregations. A single vertebrate feeding event was recorded, likely corresponding to a small gecko (Gekkonidae, possibly *Lygodactylus* sp.). Overall, animal feeding was dominated by small-bodied nocturnal insects, particularly moths, alongside substantial hemipteran-associated feeding. These patterns indicate reliance on abundant, easily captured arthropod prey, supplemented by insect-derived sugar resources.

### Seasonal diets

Plant diet composition differed significantly between seasons for both species (χ² tests, all *p* < 0.001), reflecting pronounced seasonal shifts in dominant food resources. In both species, dry-season diets were dominated by a small number of key plant taxa (e.g., *Terminalia seyrigii*), whereas wet-season diets were characterized by increased reliance on fruiting or flowering taxa such as *Operculycaria decaryi* and *Cloiselia carbonaria*, indicating close tracking of seasonal resource availability.

The overall proportion of plant versus animal-derived feeding (including both prey and hemipteran-associated feeding) remained broadly similar between seasons for both species (Figure 2). Seasonal differences in trophic proportions were not significant within either species (χ² tests, *p* > 0.05), indicating that seasonal dietary changes primarily involved shifts in the composition of foods consumed rather than major changes in overall trophic strategy. Nevertheless, comparisons between species revealed consistent trophic differentiation: across all observations, *M. murinus* consumed a significantly greater proportion of animal-derived resources than *M. griseorufus* (binomial GLM, χ² = 20.94, *p* < 0.001), whereas no overall seasonal effect was detected (χ² = 0.42, *p* = 0.516) and the Species × Season interaction was not significant (χ² = 3.02, *p* = 0.082).

Plant feeding categories showed pronounced seasonal variation (Figure 2). Across both species combined, plant-part use differed significantly between seasons (χ² = 104.01, df = 3, p < 0.001), with significant shifts detected within both species (*M. griseorufus*: χ² = 28.97, p < 0.001; *M. murinus*: χ² = 90.62, p < 0.001). During the dry season, diets were dominated by exudates (59.7% in *M. griseorufus*; 50.5% in *M. murinus*). Exudate use declined during the wet season (41.8% and 24.3%, respectively), accompanied by increased consumption of fruit and flowers, particularly in *M. murinus*, which exhibited a pronounced increase in floral feeding. However, habitat-related differences in plant-part use were comparatively modest overall, despite broader site-level differences in resource composition. Exudate use did not differ significantly between sites (χ² = 0.42, p = 0.516) and remained similar across sites in *M. griseorufus* (Mangatsiaka: 49.1%; Tsimelahy: 53.8%). In contrast, *M. murinus* showed greater habitat-related variation, with higher floral feeding at Mangatsiaka (28.6%) and fruit-dominated diets at Tsimelahy (51.5%). Multinomial logistic regression confirmed that plant-part use was significantly influenced by species (LR χ² = 21.56, p = 0.010), season (LR χ² = 157.34, p < 0.001), and site (LR χ² = 94.95, p < 0.001), with no significant Species × Season interaction (LR χ² = 4.22, p = 0.239).

Hemipteran-associated feeding showed pronounced seasonal and spatial variation. Across both species combined, the proportion of hemipteran-associated feeding increased significantly from the dry season (13.8%) to the wet season (31.7%) (χ² = 20.44, p < 0.001), although the magnitude of this shift was strongly species-dependent. In *M. griseorufus*, hemipteran-associated feeding remained low and did not differ significantly between seasons (χ² = 0.02, p = 0.902), whereas *M. murinus* showed a marked increase from 15.5% in the dry season to 46.1% in the wet season (χ² = 29.57, p < 0.001). Hemipteran-associated feeding also differed strongly between sites, being almost absent at Mangatsiaka (1.4%) but constituting a substantial component of animal-derived feeding at Tsimelahy (42.2%), with significant site effects in both species (*M. griseorufus*: χ² = 24.83, p < 0.001; *M. murinus*: χ² = 76.27, p < 0.001).

### Fecal sample analysis

Dietary remains were identified in 56 fecal samples collected across both seasons (28 per season). Plant material (including fruit, seeds, flowers, and exudates) was identified in 21 dry-season samples (75.0%) and 16 wet-season samples (57.1%). A total of fifteen seed morphotypes were identified in dry-season samples and eight in wet-season samples (Supplementary Tables S2A–B). Fruit tissues or seeds occurred in 48.2% of samples overall (15 dry-season and 12 wet-season samples), while exudate residues were detected in 14.3% of samples (four per season). Invertebrate fragments were recovered from 69.6% of samples, comprising 15 of 28 dry-season samples (53.6%) and 24 of 28 wet-season samples (85.7%). Overall, fecal analyses confirmed that both plant and animal resources contributed substantially to the diets of the study populations, while highlighting seasonal differences in the representation of dietary components, with increased prevalence of invertebrate remains in the wet season and greater representation of seeds in the dry season.

Mean recovered seed size ranged from approximately 1.4 to 8.3 mm (n = 146; median = 3.4 mm), with 64.7% of seeds measuring <5 mm, indicating dominance of small-seeded plant species consistent with dispersal of small-fruited taxa. Owing to the limited sample size, analyses were restricted to descriptive summaries, and potential species- and site-level differences were not subjected to formal statistical testing.

### Forest composition and structure

Tree community composition differed between the two forest types. The spiny thicket site (Mangatsiaka; 2,753 trees) included 111 species, whereas the transitional forest (Tsimelahy; 3,356 trees) included 126 species. Species accumulation curves approached but did not reach an asymptote, indicating incomplete sampling. Richness estimators suggested that true species richness likely exceeds observed values, with Chao1 projecting approximately 139 tree species at Mangatsiaka and 168 at Tsimelahy.

Species diversity was moderate overall (Shannon = 3.45; evenness = 0.74), but higher in the transitional forest (Shannon = 3.70, Simpson = 0.95, evenness = 0.76) than in the spiny thicket (Shannon = 3.10, Simpson = 0.91, evenness = 0.66), indicating a richer and more evenly distributed community at Tsimelahy. Trees in the spiny thicket were significantly taller on average (mean ± SD: 5.01 ± 2.64 m) than those in the transitional forest (3.35 ± 1.47 m; Welch t-test and Wilcoxon test, p < 0.0001), reflecting structural differences between vegetation types. Overall, Mangatsiaka was characterized by lower compositional heterogeneity and dominance of xerophytic taxa, whereas Tsimelahy supported a more heterogeneous assemblage including both dry-adapted and mesic species. These compositional and structural differences align with the contrasting phenological patterns observed between sites.

### Phenology

Phenological activity showed strong seasonal differences at both study sites but differed in magnitude between forest types (Table 2). In the spiny thicket forest of Mangatsiaka, only 2.3% of 2,753 trees exhibited any phenological activity (flowering, fruiting, or leaf flush) during the dry season, whereas 97.4% were inactive. Flowering (0.3%), fruiting (0.8%), and new leaf production (1.3%) were all rare during this period. In contrast, phenological activity increased markedly in the wet season, with 74.2% of 2,322 trees active and 25.5% inactive. New leaf production was the dominant phenophase (54.5%), followed by fruiting (17.6%) and flowering (10.9%).

**Table 2.** Seasonal variation in vegetation phenology at Mangatsiaka and Tsimelahy, Andohahela National Park.

| Site | Season | No. Trees | Richness | Active (%) | Inactive (%) | Flowering (%) | Fruiting (%) | New leaves (%) |
| --- | --- | --- | --- | --- | --- | --- | --- | --- |
| Mangatsiaka (spiny thicket) | Dry | 2753 | 115 | 2.3 | 97.4 | 0.3 | 0.8 | 1.3 |
|  | Wet | 2322 | 107 | 74.2 | 25.5 | 10.9 | 17.6 | 54.5 |
| Tsimelahy (transitional forest) | Dry | 3339 | 129 | 7.5 | 92.5 | 0.7 | 6.5 | 0.3 |
|  | Wet | 3299 | 125 | 50.3 | 49.7 | 10.6 | 8.7 | 34.5 |

Vegetation phenology summaries across the two study sites (Mangatsiaka, spiny thicket; Tsimelahy, transitional forest) and seasons (dry and wet). Phenological activity is expressed as the percentage of trees classified as active (showing any reproductive or vegetative activity) or inactive, and the percentage of individuals exhibiting flowering, fruiting, or new leaf production. Percentages are calculated relative to the total number of trees surveyed at each site and season.

In the transitional forest of Tsimelahy, phenological activity was also highly seasonal but less extreme. During the dry season, 7.5% of 3,339 trees were active and 92.5% inactive, with fruiting (6.5%) more common than flowering (0.7%) or new leaf production (0.3%). In the wet season, 50.3% of 3,299 trees were phenologically active and 49.7% inactive, with new leaf production again dominant (34.5%), followed by fruiting (8.7%) and flowering (10.6%). Overall, both forests exhibited pronounced seasonal shifts, with the strongest contrast observed in the spiny thicket forest, indicating stronger seasonal suppression of phenological activity than in the transitional forest.

### Seasonal resources

Despite the near-complete suppression of phenological activity during the dry season, a small subset of tree species remained active and represented important dry-season resources for lemurs. In the spiny thicket forest of Mangatsiaka, only 24 tree species exhibited any dry-season activity (flowering, fruiting, or leaf flush). These events were rare and taxonomically dispersed, with most species represented by one or two individuals (Supplementary Table S3A). The most frequently active taxa were Katrafay (*Cedrelopsis grevei*; Ptaeroxylaceae) and Hazomby (*Indigofera* sp.; Fabaceae), which showed limited flowering and leaf flush, and Maintyfo (*Diospyros* sp.; Ebenaceae), which exhibited fruiting during the dry season. A small number of additional species showed only isolated activity.

In the transitional forest of Tsimelahy, a greater number of species (31) exhibited dry-season phenological activity, most often fruiting (Supplementary Table S3B). Activity was strongly dominated by a small number of taxa, particularly Avoha (*Dichrostachys humbertii*; Mimosaceae), *Indigofera* sp., and Daromamy (*Commiphora* sp.; Burseraceae), which together accounted for a substantial proportion of fruiting records. Overall, the transitional forest retained a broader and more fruit-biased pool of dry-season resources than the spiny thicket forest, consistent with the weaker seasonal bottleneck observed at Tsimelahy. The rarity and low abundance of active species in both forests highlight the severity of seasonal resource scarcity in these dry forests. Notably, only a subset of these species appeared to contribute directly to lemur diets, with fruiting taxa such as Andranoty (*Rothmannia* sp.; Rubiaceae) and Matsaky (*Tarenna pruinosum*; Rubiaceae) representing important dry-season food resources used by both species.

## Discussion

Our results demonstrate that sympatric *Microcebus murinus* and *M. griseorufus* at Andohahela National Park exploit a broadly similar, plant-dominated resource base, yet exhibit consistent differences in key aspects of their feeding ecologies. Both species relied heavily on plant resources, consuming a wide diversity of shared taxa at different proportions across the year. *Microcebus murinus* consistently consumed a greater proportion of animal-derived resources across seasons and showed marked increases in floral feeding and hemipteran-associated secretion feeding during the wet season. In contrast, *M. griseorufus* relied more strongly on exudates throughout the year, consistent with previous studies highlighting the importance of exudates in the diet of this species (Génin, 2001). Our findings indicate that, despite substantial overlap in the types of resources exploited, the two species occupy subtly differentiated trophic niches within the flexible, omnivorous ecological framework characteristic of mouse lemurs (Martin, 1972; Hladik, 1979; Richard & Dewar, 1991; Atsalis, 1999; Ganzhorn et al., 2023).

### Plant resource use and dietary divergence

Plant resource use in this system is structured along a gradient of temporal predictability, ranging from ephemeral floral and fruit resources to stable exudates, with marked interspecific differences in how these resources are exploited (Andriaharimalala et al., 2012).

### Fruit as a shared and opportunistic resource

Fruit constituted a substantial component of the plant diet in both species and increased during the wet season as exudate use declined, reflecting seasonal changes in phenological cycles and resource availability. Although fruit availability shows a clear seasonal pulse, particularly in the spiny thicket habitat, some fruit resources remain available across seasons relative to more ephemeral plant foods. Consistent with this, fruit consumption was broadly similar between species (36–37% of identified plant feeding records), indicating that fruit represents a shared and widely exploited resource rather than a major axis of dietary differentiation. Fruit use was structured around a limited number of taxa in both species but showed weaker dominance patterns than exudate or floral feeding. In *M. griseorufus*, wet-season fruit use was strongly dominated by Jahiby (41.7%), indicating local specialization on a highly abundant resource. In contrast, *M. murinus* exhibited a more distributed pattern, with multiple co-dominant taxa contributing substantially in both seasons (including Jahiby; 30.6% in the wet season). This more even distribution suggests a greater tendency toward multi-resource exploitation rather than reliance on a single dominant fruit taxon.

Comparable patterns have been reported across mouse lemur systems, where fruit consumption increases during periods of high availability. In northwestern Madagascar, for example, *M. murinus* can shift toward strongly fruit-dominated diets during the wet season, with fruit comprising up to ∼80% of feeding records (Thorén et al., 2011). Similarly, high fruit consumption has been reported in humid forests species such as *M. rufus* (Atsalis, 1999) and *M. tanosi* (unpublished data), indicating that reliance on fruit is widespread, though variable, across the genus. In contrast, in other sympatric systems, *M. ravelobensis* tends to maintain more mixed diets, with fruits comprising lower dietary contributions (Thorén et al., 2011), while at Kirindy fruits and flowers contribute relatively little to the diet of *M. berthae* (<10%; Dammhahn & Kappeler, 2008a). Taken together, these findings indicate that fruit is a broadly accessible and opportunistically exploited resource that contributes to seasonal dietary shifts but plays a limited role in interspecific dietary divergence.

### Floral resources and pulse tracking in *M. murinus*

Floral resources represented a distinct and important component of the plant diet and formed a key axis of interspecific differentiation. Consistent with underlying phenological dynamics, *Microcebus murinus* consumed flowers and nectar substantially more frequently than *M. griseorufus*, particularly during the wet season when floral resources were most abundant. Floral feeding was strongly concentrated on a small number of plant taxa, with a limited set of species contributing disproportionately to the diet, indicating targeted exploitation of key nectar resources rather than diffuse use across the plant community. Nectar is characterized by high concentrations of simple sugars and rapid accessibility, making it an energetically valuable but temporally constrained resource. The pronounced wet-season increase in floral feeding observed in *M. murinus* during the wet season therefore suggests active exploitation of short-lived carbohydrate pulses. In contrast, the more limited use of floral resources by *M. griseorufus* indicates reduced reliance on these ephemeral energy sources, consistent with its greater dependence on exudates. These differences reinforce a broader pattern of dietary divergence in this system, suggesting that *M. murinus* more strongly exploits temporally variable, high-reward resources, while *M. griseorufus* maintains a more stable feeding strategy centered on predictable plant-derived foods.

### Exudates as predictable resources

Exudates formed a central component of the plant diet in both species, particularly during the dry season, highlighting their importance as reliable resources in highly seasonal environments. They were consistently more important in the diet of *M. griseorufus* than in *M. murinus*, especially during the dry season when exudates dominated plant feeding (approximately 60% of records in *M. griseorufus* and 50% in *M. murinus*). Although their contribution declined in the wet season, exudates remained a substantial component of the diet, particularly in *M. griseorufus*. Exudate use was strongly concentrated on a small number of plant taxa, dominated by *Terminalia seyrigii*, with secondary contributions from *Commiphora aprevalii* and *Operculycaria decaryi*. The prominence of *T. seyrigii* likely reflects both its nutritional value and its availability, as it is among the more common tree species at both study sites. Despite broad taxonomic dietary breadth, exudate feeding was therefore structured around a limited set of predictable, high-value resources.

Similar patterns have been documented elsewhere, particularly where *Terminalia* exudates form a major dietary component during the dry season (Radespiel et al., 2006). In northwestern Madagascar, for example, gum constitutes a substantial proportion of the diet of *Microcebus murinus*, dominating feeding records during the dry season and remaining important outside peak fruiting periods (Thorén et al., 2011; Radespiel et al., 2006). In the same system, sympatric *M. ravelobensis* maintains a more mixed feeding strategy but still relies heavily on exudates, which consistently contribute a large proportion of the diet across seasons (Thorén et al., 2011). These gums are rich in soluble sugars and contain moderate levels of protein, providing a valuable resource when other foods are scarce (Hladik et al., 1980; Nash, 1986). However, the extent of exudate use varies among systems. For example, in Kirindy, exudates contribute relatively little to the diet of *M. murinus* and are rarely used by *M. berthae* (Dammhahn & Kappeler, 2008a), suggesting strong ecological context dependence. Exudates are available year-round and are more spatially and temporally predictable than fruit or floral resources. As such, they represent a low-risk foraging strategy that can buffer individuals against environmental variability. The stronger reliance on exudates in *M. griseorufus* is consistent with a feeding strategy centered on predictable, structurally accessible plant-derived resources, whereas the reduced reliance in *M. murinus* reflects greater exploitation of temporally variable, high-reward foods.

### Animal resources

Animal-derived feeding constituted a substantial component of the diet in both species but was consistently more pronounced in *Microcebus murinus*. True prey records were dominated by small, abundant invertebrate prey, particularly Lepidoptera, indicating reliance on readily available and energetically efficient prey types. Similar patterns have been reported in other mouse lemur systems, where arthropod feeding is often dominated by common taxa such as Coleoptera and Lepidoptera, with relatively little evidence for strong taxonomic specialization in prey capture (Hladik et al., 1980; Atsalis, 1999; Dammhahn & Kappeler, 2008a). When hemipteran-associated feeding is excluded, both species consumed comparable proportions of insect prey, indicating that interspecific differences in animal-derived feeding are driven primarily by differential use of hemipteran-associated carbohydrate resources rather than differences in prey capture. Proportions of true prey were also broadly similar across seasons in both species, indicating that seasonal variation in animal-derived feeding is driven primarily by shifts in hemipteran-associated resource use rather than changes in prey consumption. These patterns are consistent with those reported for the sympatric *M. murinus–M. berthae* system in Kirindy, where both species consumed similar proportions and types of arthropod prey despite differences in overall feeding strategy (Dammhahn & Kappeler, 2008a). In contrast, hemipteran records in this system predominantly reflect hemipteran-associated honeydew feeding rather than active prey capture, with only a single cicada record representing true predation. This distinction highlights the dual role of arthropods in mouse lemur diets, functioning both as prey and as sources of secondary carbohydrate resources.

Hemipteran-associated feeding represented a substantial and functionally distinct component of animal-derived feeding. Across Madagascar, sap-feeding fulgoroid insects, including flatids, are known to provide indirect carbohydrate resources via honeydew and related secretions that may be consumed directly or as residues deposited on vegetation (Hladik et al., 1980; Corbin & Schmid, 1995; Radespiel et al., 2006; Dammhahn & Kappeler, 2008a). In our system, the use of these resources differed markedly between species, seasons, and sites. Across both species combined, the proportion of animal-derived feeding records attributable to hemipteran-associated feeding increased from 13.8% in the dry season to 31.7% in the wet season. However, this pattern was strongly species-dependent: *M. griseorufus* showed little seasonal change (11.1% in the dry season; 12.7% in the wet season), whereas *M. murinus* exhibited a marked increase from 15.5% to 46.1%. Hemipteran-associated feeding also differed strongly between sites, being almost absent at Mangatsiaka (1.4%) but forming a substantial component of animal-derived feeding at Tsimelahy (42.2%). The strong seasonal increase in hemipteran-associated feeding in *M. murinus* likely reflects variation in resource availability. Honeydew production by sap-feeding hemipterans is closely linked to plant productivity and insect abundance, both of which decline under dry-season conditions. This interpretation is further supported by its strong spatial concentration within the more productive transitional forest habitat.

These results indicate that *M. murinus* exhibits pronounced plasticity in its use of hemipteran-associated resources, particularly under wet-season and transitional-forest conditions. In contrast, *M. griseorufus* maintains a more conservative pattern, with comparatively limited use of hemipteran-associated feeding and continued reliance on other insect prey types. This divergence is consistent with the broader dietary contrast observed throughout the study, whereby *M. murinus* more strongly exploits temporally and spatially variable sugar-rich resources, while *M. griseorufus* relies less on these ephemeral carbohydrate sources and more on predictable plant-derived foods.

The increase in hemipteran-associated feeding observed in *M. murinus* is consistent with patterns reported from Kirindy (Dammhahn & Kappeler, 2008a), but appears particularly pronounced in the present study. This difference may reflect variation in interspecific competition for secretion resources across sympatric systems. In Kirindy, hemipteran secretions are heavily exploited by both species, but particularly by *M. berthae*, which can devote up to 90% of its feeding time to this resource during the dry season (Dammhahn & Kappeler, 2008a), whereas in our system *M. griseorufus* shows comparatively limited use of hemipteran-associated resources. Reduced competition for this resource may therefore help explain the stronger shift observed in *M. murinus*. At Ankarafantsika, by contrast, sympatric *M. murinus* and *M. ravelobensis* both make use of insect secretions, although reliance on this resource appears to be higher and more consistent in *M. ravelobensis* (Thorén et al., 2011), indicating a shared but asymmetrical dependence on hemipteran-associated carbohydrate resources. Taken together, these systems indicate that sympatric mouse lemur assemblages span a continuum in the use of hemipteran-associated resources, from extreme reliance in *M. berthae*, to moderate use in *M. ravelobensis*, to comparatively limited use in *M. griseorufus*, with *M. murinus* flexibly adjusting its use of this resource according to ecological context.

### Fecal evidence and dietary representation

Fecal analyses provided complementary insights into dietary composition but revealed patterns that differed from direct behavioral observations, with a higher prevalence of invertebrate remains in wet-season samples and increased representation of seeds in dry-season samples. These differences likely reflect a combination of sampling artifacts, seasonal shifts in resource use, and differential detectability of dietary components in fecal material. Seeds and chitinous insect fragments are readily recovered after digestion, whereas nectar, exudates, and hemipteran-associated secretions—despite forming important components of the diet—are less likely to leave identifiable traces.

The greater occurrence of seeds in dry-season samples appears to be driven in part by *M. murinus*, suggesting continued exploitation of scarce fruit resources by this species even during periods of reduced overall availability. In contrast, seeds were relatively infrequent in *M. griseorufus* samples, consistent with its stronger reliance on exudates, which are poorly represented in fecal material. Notably, tree sap residues were more frequently detected in *M. murinus* samples despite observational evidence of greater exudate use by *M. griseorufus*. This discrepancy likely reflects differences in detectability and processing of exudates, which may leave limited residues in fecal material. Mean recovered seed sizes overlapped broadly between species, with most seeds falling within the small-size range (<5 mm). Conversely, the higher prevalence of invertebrate remains (i.e., true prey) in wet-season samples is consistent with increased insect availability (Wolda, 1988; Dammhahn & Kappeler, 2008a) and opportunistic foraging during periods of elevated productivity.

### Synthesis: carbohydrate strategies and energy storage

Together, these patterns indicate that resource use is structured along a gradient from temporally variable, high-reward resources to predictable, low-variance resources, with *M. murinus* and *M. griseorufus* occupying different positions along this axis. *M. murinus* appears to operate at the more flexible end of the gradient, tracking seasonal pulses in fruit, floral, and insect-derived resources, whereas *M. griseorufus* relies more heavily on stable plant-derived foods, particularly exudates. Rather than functioning solely as fallback foods, exudates in this system appear to represent predictable, low-variance resources that can buffer individuals against environmental unpredictability. The stronger reliance on exudates in *M. griseorufus* is therefore consistent with a strategy focused on reducing foraging risk and maintaining energy balance under variable conditions, rather than maximizing short-term energetic gain. This suggests that dietary divergence in this system reflects not only differences in resource use, but contrasting strategies for coping with environmental variability. Thus, coexistence is facilitated by fine-scale and often context-dependent differences in resource use within a broadly shared dietary framework, rather than by strict trophic partitioning.

These differences may also reflect underlying physiological strategies related to energy storage. Experimental work in the fat-tailed dwarf lemur (*Cheirogaleus medius*) has shown a preference for high-sugar fruits during pre-hibernation fattening (Fietz & Ganzhorn, 1999), suggesting that access to rapidly assimilable carbohydrates can play an important role in seasonal energy balance. A similar mechanism may operate in *M. murinus*. In this context, the strong reliance of *M. murinus* on sugar-rich resources such as fruit, nectar, and hemipteran-associated secretions may reflect an opportunistic strategy that maximizes energy intake during periods of high resource availability. In contrast, *M. griseorufus* appears to rely less on rapidly available carbohydrate sources and more on structurally predictable resources such as exudates, suggesting a different energetic strategy that may depend less on rapid energy acquisition and more on stable resource buffering and distinct physiological strategies (Hyde Roberts et al., 2026a). Although this hypothesis requires direct physiological validation, it offers a plausible mechanistic basis for the observed dietary divergence. More broadly, it suggests that differences in plant resource use in this system may reflect not only ecological opportunity, but also species-specific strategies of energy storage and use.

### Macroecological and evolutionary context

The high degree of overlap in dietary ecology and overall trophic composition is not surprising given the shared ancestry of these species (Poelstra et al., 2021; Van Elst et al., 2025) and the strong environmental constraints of the xeric forest habitats of Andohahela. While mouse lemurs are widely regarded as adaptable omnivores, our findings reveal consistent fine-scale ecological variation and subtle species-specific differences, mirroring patterns reported from other sympatric systems involving *M. murinus* (Radespiel et al., 2006; Dammhahn & Kappeler, 2008a; Thorén et al., 2011).

The broader ecological context provides a useful framework for interpreting these patterns, as reflected in the contrasting biogeographic distributions of the two species. *Microcebus murinus* is typically regarded as a highly flexible generalist, occupying one of the widest ranges of any mouse lemur species. It inhabits a variety of forest types, occurring in sympatry with multiple congeners across western and southern Madagascar (Schmid & Kappeler, 1994; Zimmermann et al., 1998; Rasoloarison et al., 2000; Olivieri et al., 2007; Gligor et al., 2009). In contrast, *M. griseorufus* is largely restricted to the southern dry and spiny forest formations, where it occupies climatically extreme and highly seasonal environments (Génin, 2008). These contrasting ecological and biogeographic histories may help explain the dietary differences observed here, and are consistent with the idea that species-specific ecological strategies are maintained under conditions of sympatry. More broadly, such patterns are compatible with models of Pleistocene allopatric divergence followed by range expansion and secondary contact. The wide distribution and ecological flexibility of *M. murinus* suggest that it may have expanded into habitats occupied by more specialized congeners (Schneider et al., 2010; Hapke et al., 2011), particularly under shifting climatic conditions (Beyer et al., 2020). In such systems, coexistence may be facilitated by the persistence of contrasting resource-use strategies, with locally adapted species relying on predictable resources, while *M. murinus* exploits transient resource pulses.

## Conclusion

Sympatric, closely related mouse lemurs at Andohahela exhibit broadly similar feeding ecologies but differ consistently in how key resources are exploited, particularly in the balance between exudates, floral resources, hemipteran-associated secretions, and animal prey. These differences reflect contrasting strategies of resource use, with *Microcebus murinus* exhibiting greater trophic flexibility and responsiveness to temporally variable, carbohydrate-rich resources, while *M. griseorufus* relies more heavily on predictable, low-variance plant-derived foods. Together, these findings highlight the importance of subtle trophic differentiation in facilitating coexistence in highly seasonal environments and suggest that dietary divergence may be shaped not only by ecological opportunity, but also by species-specific strategies for coping with environmental variability and managing energy balance.

## Acknowledgments

We thank the Ministry of Environment and Sustainable Development (MEDD), CAFF/CORE, Madagascar National Parks (MNP), and the Université d’Antananarivo for supporting this project over several years. We particularly thank Prof. Fanomezana Ratsoavina, Jacques Francky Rakotoarisoa, Lalatiana Odile Randriamiharisoa, Garina Parfait Manantsoa, and Dan Vick for their assistance in facilitating the research permitting process. We extend our sincere gratitude and respect to our Malagasy field team, whose knowledge, commitment, and hard work made this research possible. Finally, we wish to acknowledge the wider communities of Andohahela, whose cordial hospitality provided the backdrop for this work. This work was funded by NSFDEB-2148914 to ADY.

## Supplementary Materials

**Table S1A.**
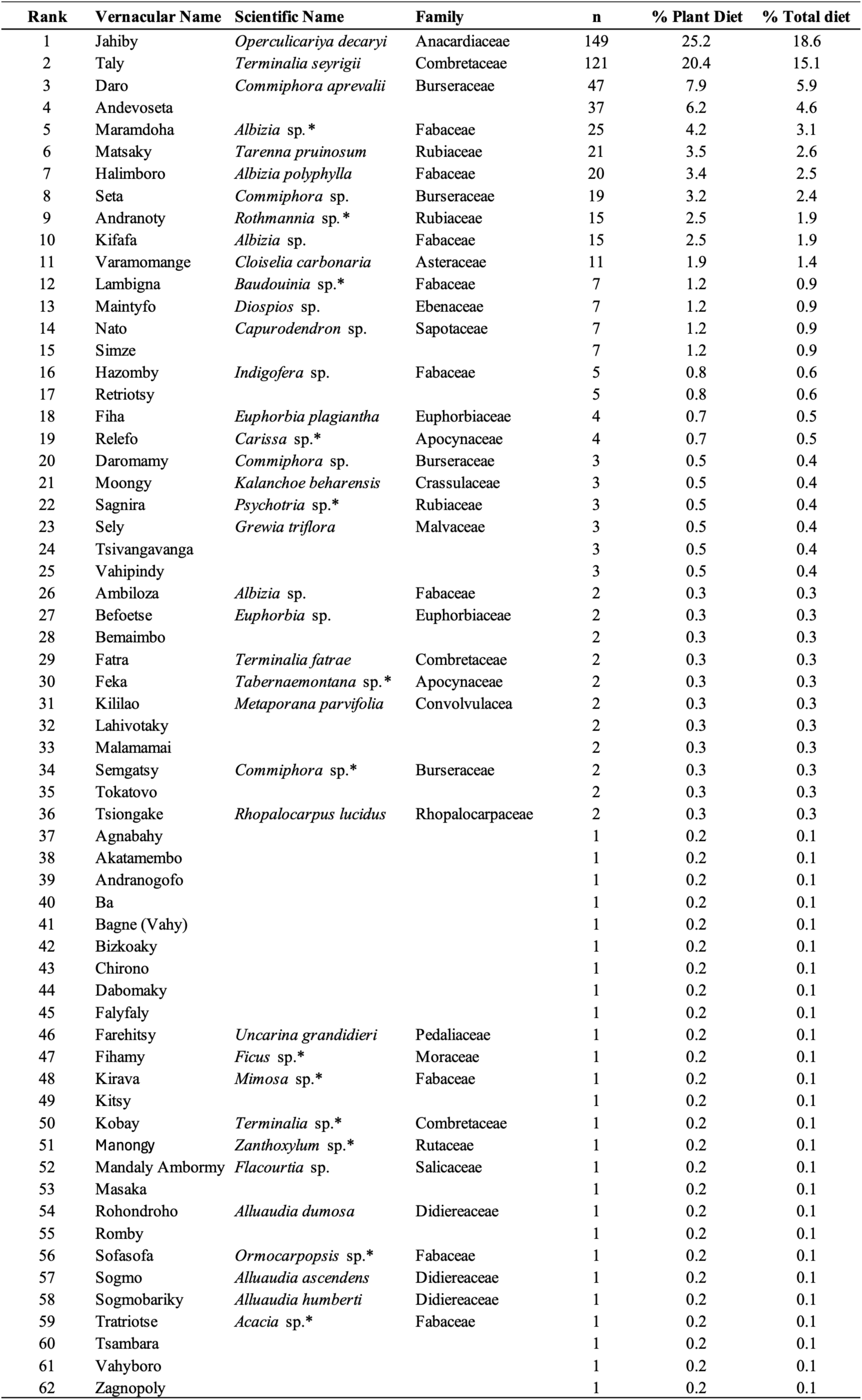
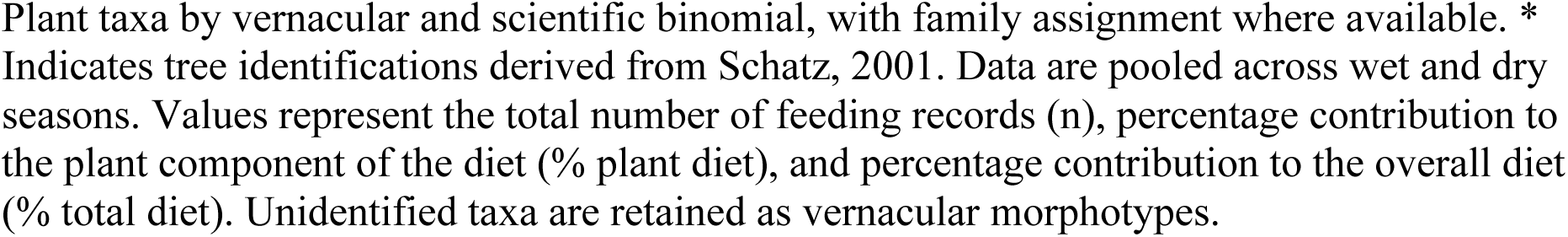
Plant taxa recorded in the diet of *Microcebus griseorufus* at Andohahela National Park.

**Table S1B.**
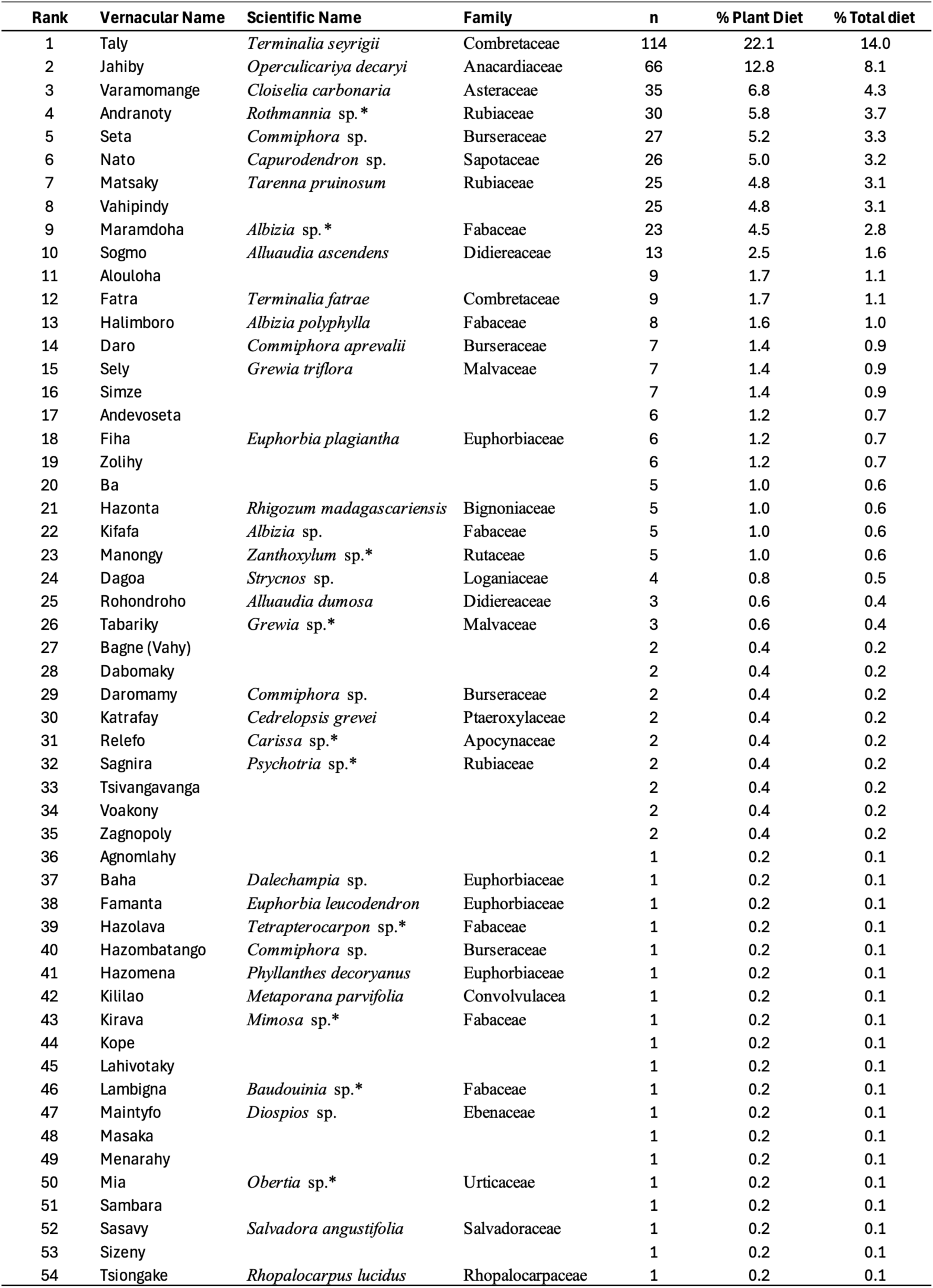
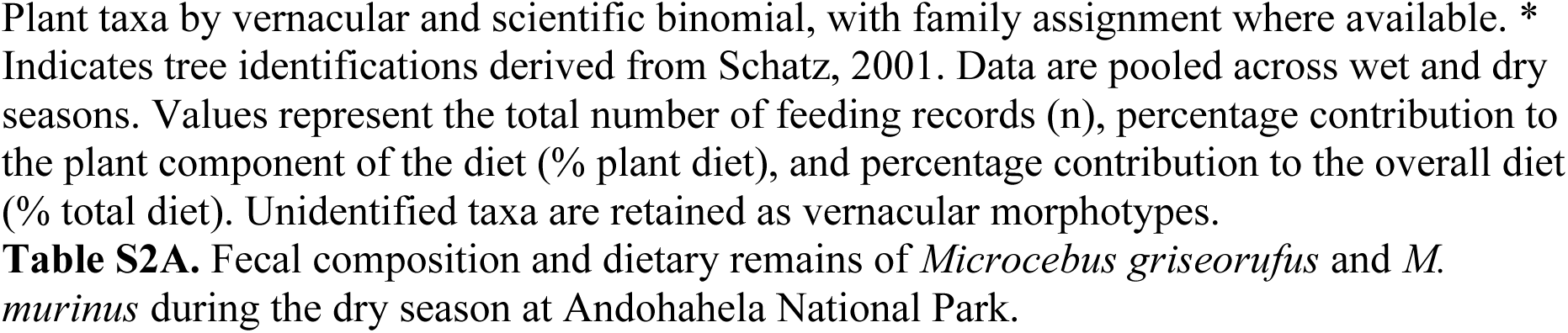
Plant taxa recorded in the diet of *Microcebus murinus* at Andohahela National Park.

**Table S2A.** Fecal composition and dietary remains of *Microcebus griseorufus* and *M. murinus* during the dry season at Andohahela National Park.

| Season | Site | Sample | Species | No. Seeds | Seed Types | Mean Size (mm) | Plant | Fruit / Seeds | Tree sap | Invertebrates |
| --- | --- | --- | --- | --- | --- | --- | --- | --- | --- | --- |
| Dry | Mangatsiaka | S1 | *M. griseorufus* |  |  |  |  |  |  | ✓ |
|  |  | S2 | *M. griseorufus* | 5 | 1 | 3.3 | ✓ | ✓ |  |  |
|  |  | S3 | *M. griseorufus* |  |  |  |  |  |  | ✓ |
|  |  | S4 | *M. murinus* | 1 | 1 | 4.6 | ✓ | ✓ |  | ✓ |
|  |  | S5 | *M. murinus* |  |  |  | ✓ |  | ✓ | ✓ |
|  |  | S6 | *M. murinus* | 1 | 1 | 5.5 | ✓ | ✓ |  | ✓ |
|  |  | S7 | *M. murinus* | 1 | 1 | 5.2 | ✓ | ✓ |  | ✓ |
|  |  | S8 | *M. murinus* |  |  |  | ✓ |  | ✓ | ✓ |
|  |  | S9 | *M. murinus* | 8 | 1 | 2.2 | ✓ | ✓ |  | ✓ |
|  |  | S10 | *M. murinus* |  |  |  |  |  |  | ✓ |
|  |  | S11 | *M. murinus* |  |  |  | ✓ |  | ✓ |  |
|  |  | S12 | *M. murinus* |  |  |  |  |  |  | ✓ |
|  | Tsimelahy | S13 | *M. griseorufus* | 4 | 1 | 5.7 | ✓ | ✓ |  | ✓ |
|  |  | S14 | *M. griseorufus* |  |  |  |  |  |  | ✓ |
|  |  | S15 | *M. griseorufus* | 1 | 1 | 5.0 | ✓ | ✓ |  |  |
|  |  | S16 | *M. griseorufus* |  |  |  |  |  |  | ✓ |
|  |  | S17 | *M. griseorufus* |  |  |  |  |  |  | ✓ |
|  |  | S18 | *M. griseorufus* |  |  |  | ✓ |  |  |  |
|  |  | S19 | *M. griseorufus* |  |  |  | ✓ |  |  |  |
|  |  | S20 | *M. griseorufus* | 1 | 2 | 7.0, 5.0 | ✓ | ✓ |  |  |
|  |  | S21 | *M. griseorufus* | 4 | 1 | 5.2 | ✓ | ✓ |  |  |
|  |  | S22 | *M. murinus* | 2 | 1 | 4 | ✓ | ✓ |  | ✓ |
|  |  | S23 | *M. murinus* | 6 | 2 | 5.2, 2.1 | ✓ | ✓ |  |  |
|  |  | S24 | *M. murinus* | 3 | 1 | 3.4 | ✓ | ✓ |  |  |
|  |  | S25 | *M. murinus* | 7 | 1 | 5.1 | ✓ | ✓ |  |  |
|  |  | S26 | *M. murinus* | 4 | 1 | 2.1 | ✓ | ✓ |  |  |
|  |  | S27 | *M. murinus* |  |  |  | ✓ |  | ✓ |  |
|  |  | S28 | *M. murinus* | 3 | 1 | 7.6 | ✓ | ✓ |  |  |

Fecal sample analysis, summarized by collection study site (Mangatsiaka and Tsimelahy) and species. Presence of dietary components is indicated by ✓. Seed morphospecies (number of distinct morphotypes) were as follows: Mangatsiaka—*M. griseorufus* = 1, *M. murinus* = 3; Tsimelahy—*M. griseorufus* = 4, *M. murinus* = 8.

**Table S2B.**
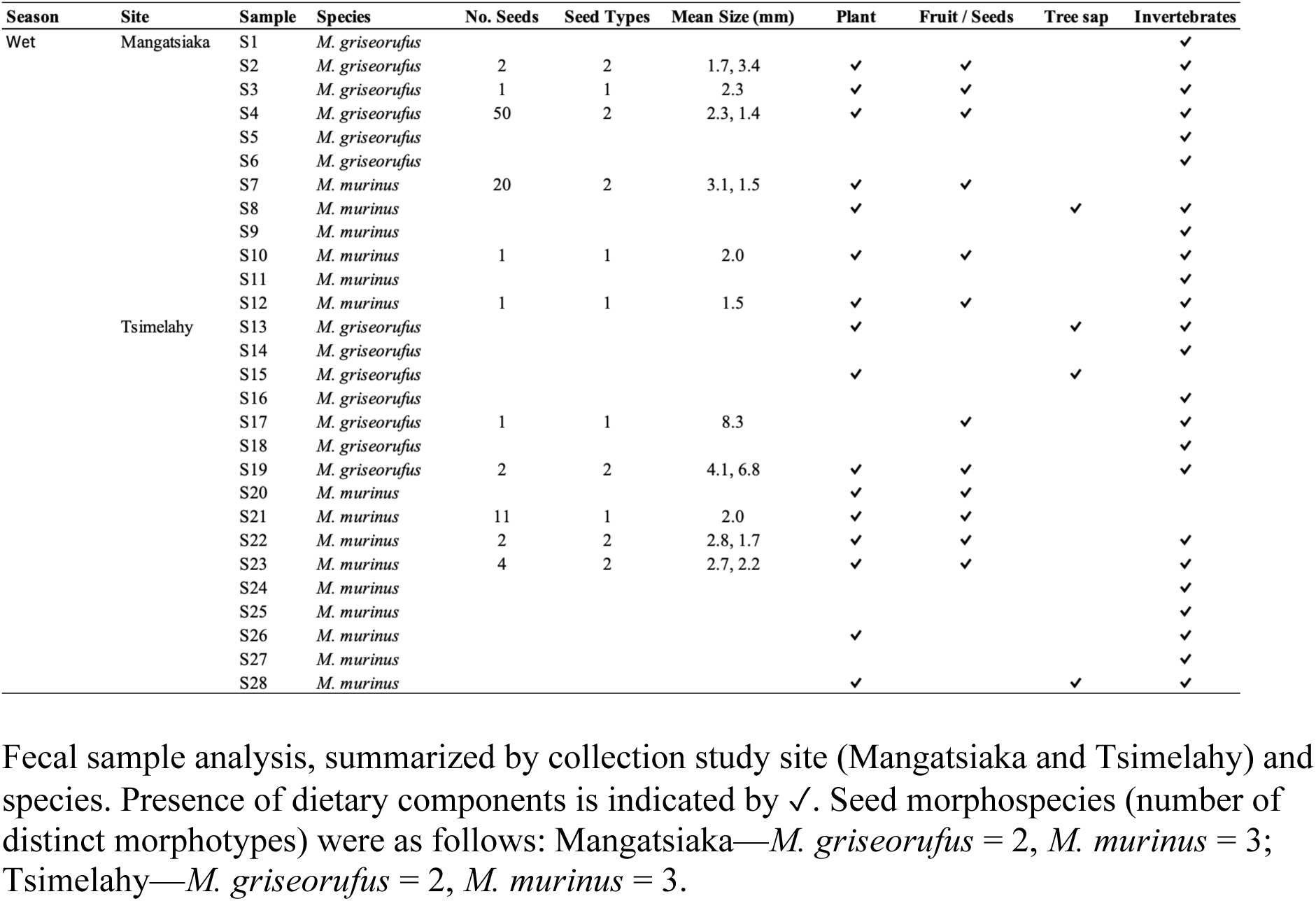
Fecal composition and dietary remains of *Microcebus griseorufus* and *M. murinus* during the wet season at Andohahela National Park.

**Table S3A.**
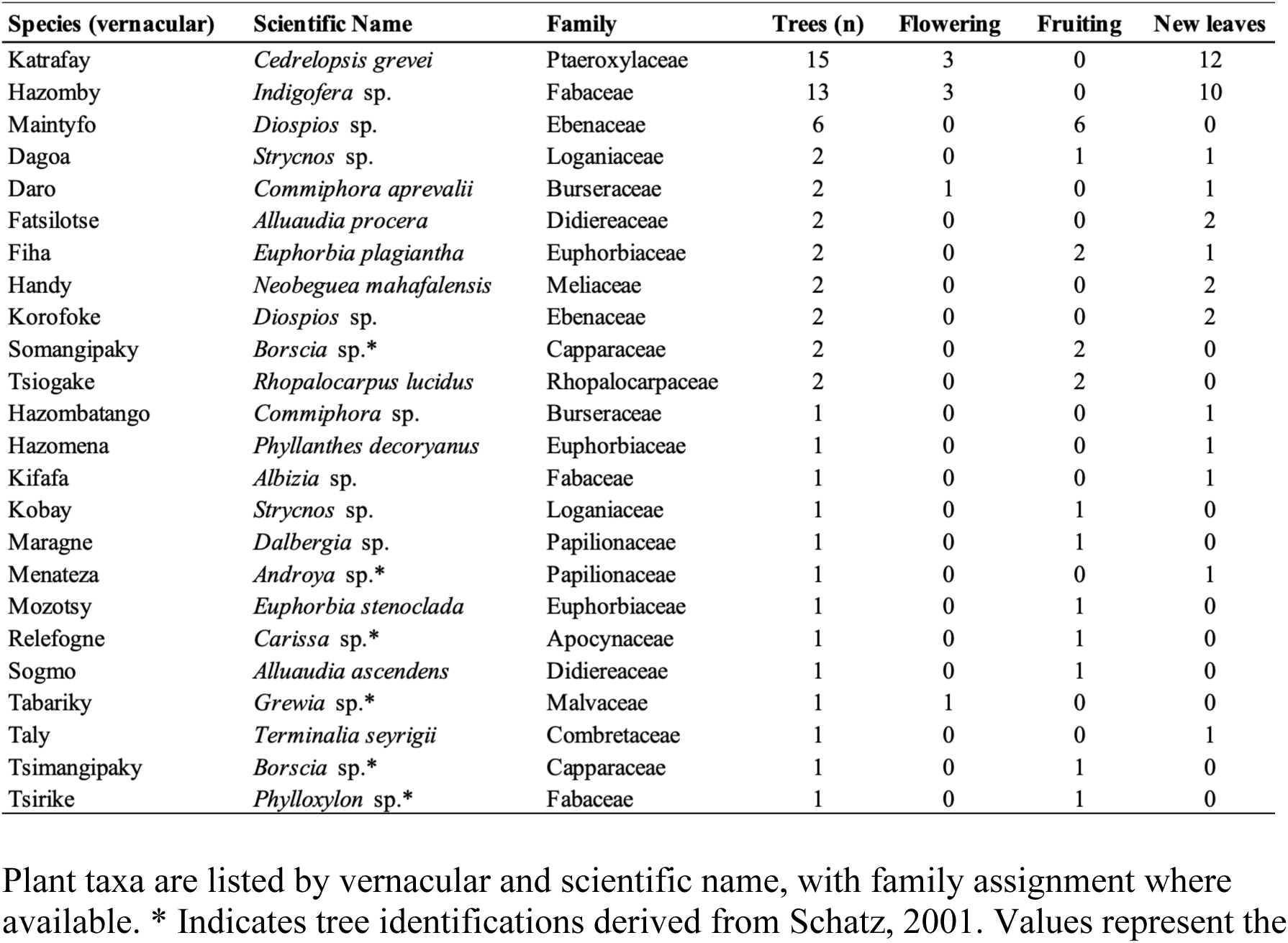
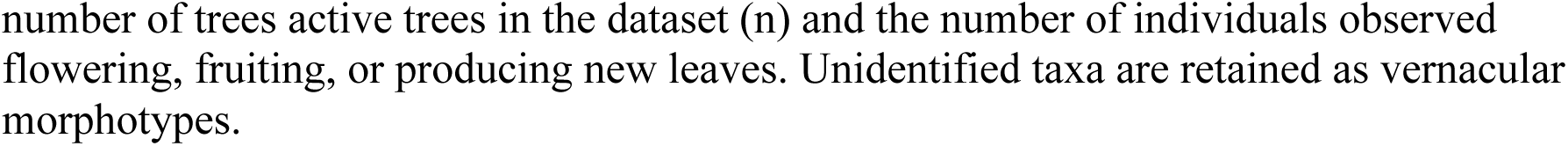
Dry-season plant resources recorded at Mangatsiaka (spiny thicket), Andohahela National Park.

**Table S3B.**
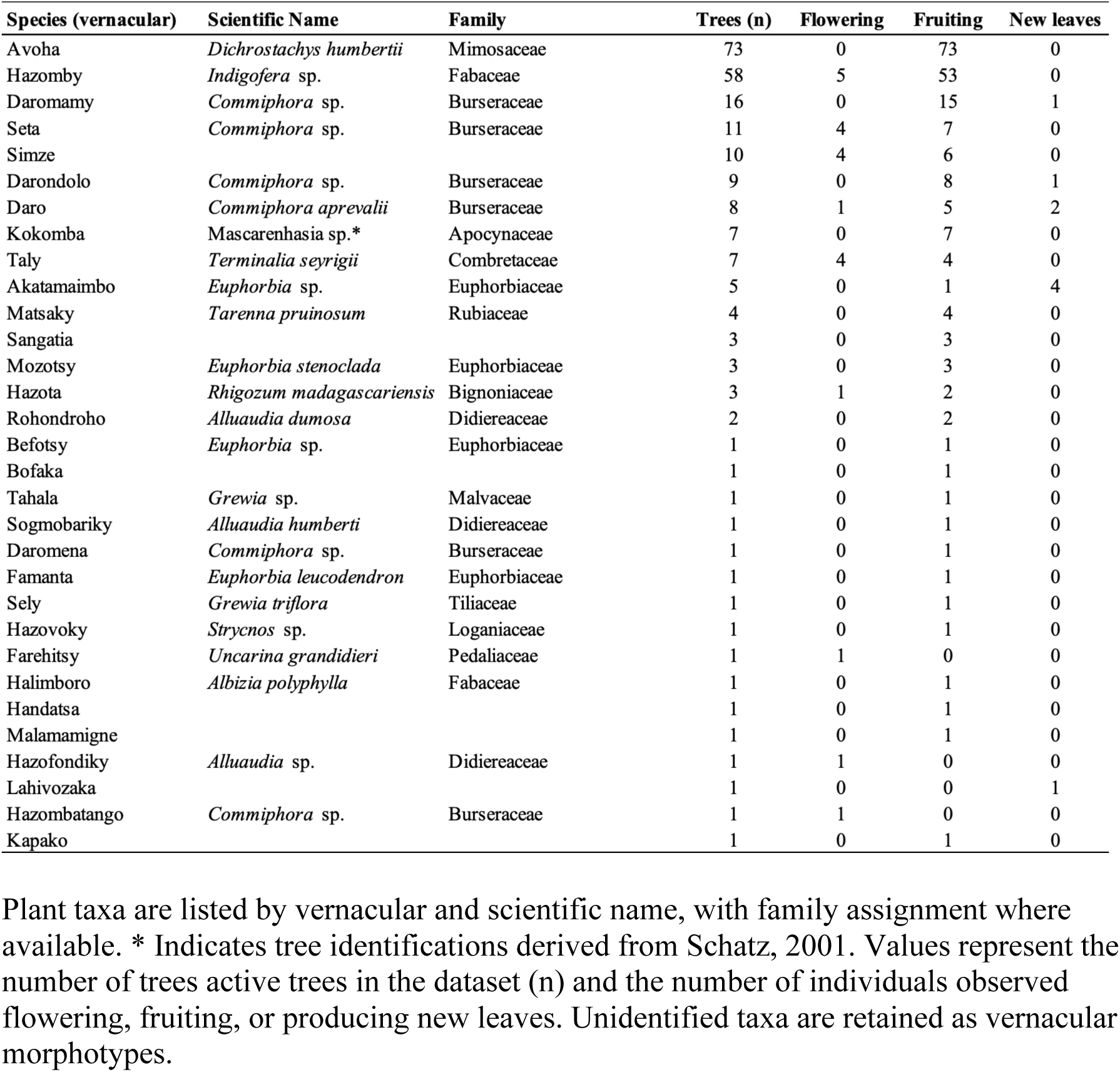
Dry-season plant resources recorded at Tsimelahy (spiny thicket), Andohahela National Park.

